# A versatile cohesion manipulation system reveals CENP-A dysfunction accelerates female reproductive age-related egg aneuploidy

**DOI:** 10.1101/2025.02.27.640570

**Authors:** Jiyeon Leem, Tom Lemonnier, Ani Khutsaidze, Lei Tian, Xiaojun Xing, Suxia Bai, Timothy Nottoli, Binyam Mogessie

**Affiliations:** Department of Molecular, Cellular and Developmental Biology, Yale University, New Haven, Connecticut, USA; Yale Genome Editing Center, Yale University School of Medicine, New Haven, Connecticut, USA; Department of Comparative Medicine, Yale Genome Editing Center, Yale University School of Medicine, New Haven, Connecticut, USA; Department of Obstetrics, Gynecology and Reproductive Sciences, Yale University School of Medicine, New Haven, Connecticut, USA

## Abstract

Female reproductive aging is accompanied by a dramatic rise in the incidence of egg aneuploidy. Premature loss of chromosome cohesion proteins and untimely separation of chromosomes is thought to underly high rates egg aneuploidy during maternal aging. However, because chromosome cohesion loss occurs gradually over female reproductive lifespan and cytoskeletal defects alone can predispose eggs to chromosomal abnormalities, the root causes of exponential rise in egg aneuploidy at advanced reproductive ages remain a mystery. Here, we applied high-resolution live imaging to visualize for the first time cohesion protein dynamics underpinning meiotic chromosome segregation. To discover proteins whose dysfunction accelerates aneuploidies associated with female reproductive aging, we innovated the first experimental system in which chemically induced cohesion reduction rapidly triggers aging-like chromosomal abnormalities in young eggs. By integrating this direct cohesion manipulation system with quantitative high-resolution microscopy and targeted protein degradation tools, we identified the centromeric protein CENP-A as a new factor whose aging-like depletion causes a dramatic rise in premature separation of sister chromatids. Our work illuminates cohesion loss-independent origins of age-related egg aneuploidy and provides new avenues to discover therapeutic targets for extending the female reproductive lifespan.

## Main text

Chromosome segregation is critical for eukaryotic cell division. When fertilizable eggs are formed from progenitor oocytes, a specialized form of meiotic cell division separates the chromosomes. Accurate chromosome segregation in mammalian oocytes and eggs is driven by a meiotic spindle machinery that is assembled from microtubules (*1*) and is reinforced with spindle actin filaments that mediate chromosome-microtubule attachments (*2, 3*). Unlike somatic cell mitosis or sperm meiosis, female meiotic chromosome segregation is remarkably susceptible to errors (*4, 5*). Egg aneuploidy, a cellular state of incorrect chromosome numbers, that arises from oocyte chromosome segregation errors typically leads to chromosomal abnormalities associated with miscarriages, infertility and genetic conditions including intellectual disabilities (e.g. Down syndrome) (*4–6*). Importantly, the incidence of egg aneuploidy increases almost exponentially at advanced female reproductive ages (*4–6*), which dramatically impacts the quality of eggs and leads to poor reproductive outcomes. With a record high maternal age at first pregnancy (*7–10*), declining fertility and a rapidly aging global human population (*11–16*), uncovering the molecular origins of female reproductive age-related egg aneuploidy is perhaps one of the most urgent clinical and socioeconomic challenges of our era (*9*).

Female reproductive age-related loss of chromosome cohesion is thought to underly high chromosome segregation error rates in oocytes (*4–6*). Consistently, in mice (*17–19*) and humans (*20*), REC8, SGO2 and other cohesin components gradually become depleted with advancing maternal age. However, given the progressive nature of cohesion weakening as well as actin and microtubule cytoskeletal defects that are common in aged oocytes (*2, 21*), it is conceivable that cohesion depletion-independent causes of chromosome segregation errors trigger the sharp rise in oocyte aneuploidy during aging. The lack of tools for direct and temporally resolved manipulation of chromosome cohesion in oocytes has remained a major hurdle in uncovering these root causes of age-related oocyte aneuploidy.

We reasoned that discovering novel causes of age-related egg aneuploidy necessitates tools for controlled generation of aging-like premature chromatid separation events in young oocytes. To achieve this, we innovated a versatile experimental system that enables direct observation of REC8, a meiosis-specific cohesin subunit (*22, 23*), in live oocytes and allows its destruction via <u>pro</u>teolysis <u>ta</u>rgeting <u>c</u>himera (PROTAC) (*24–26*) or single variable domain antibody (nanobody) (*27–29*) technologies.

We previously showed that the dTAG system, a protein degradation tool wherein the PROTAC dTAG-13 targets FKBP12^F36V^-chimeric proteins for proteasomal degradation (*30, 31*) (Fig. S1A), can successfully be used to rapidly degrade exogenously expressed actin mutants in mammalian oocytes (*32*). In addition, we demonstrated that the TRIM-Away method of rapid protein degradation can be coupled with GFP nanobodies for efficient removal of fluorescence-tagged endogenous proteins (*33*). To extend these approaches to targeted degradation of chromosome cohesion proteins and generation of prematurely separated chromatids, we applied CRISPR knockin technology and engineered mice in which REC8 is C-terminally tagged with FKBP12^F36V^-mClover3 (Fig. S1, B and C).

We validated that this endogenous protein tagging approach did not interfere with REC8 protein localization and function by immunostaining homozygous REC8-FKBP12^F36V^-mClover3 mouse oocytes with anti-GFP antibodies that cross-react with mClover3 (Fig. S1D). Three-dimensional high-resolution immunofluorescence microscopy assays demonstrated that anti-GFP antibodies recognize REC8 protein along the arms and centromeric regions of homologous chromosomes (Figs. 1A and S1E; Movie S1), confirming that fusion to FKBP12^F36V^-mClover3 does not impact its localization. Sequential removal of REC8 cohesin from chromosome arms in meiosis I and from centromeric regions of DNA in meiosis II underpins successive separation of homologous chromosomes and sister chromatids (*5, 34, 35*). High-spatiotemporal resolution microscopy of SiR-5-Hoechst labelled chromosomes and REC8-FKBP12^F36V^-mClover3 allowed us for the first time to directly visualize this selective REC8 degradation phenomenon in live oocytes undergoing meiosis I chromosome segregation (Fig. 1B; Movies S2). Consistent with our immunofluorescence microscopy data (Fig. 1A), these observations confirmed that oocyte REC8 cohesin dynamics is unaffected in these engineered mice.

**Fig. 1.**
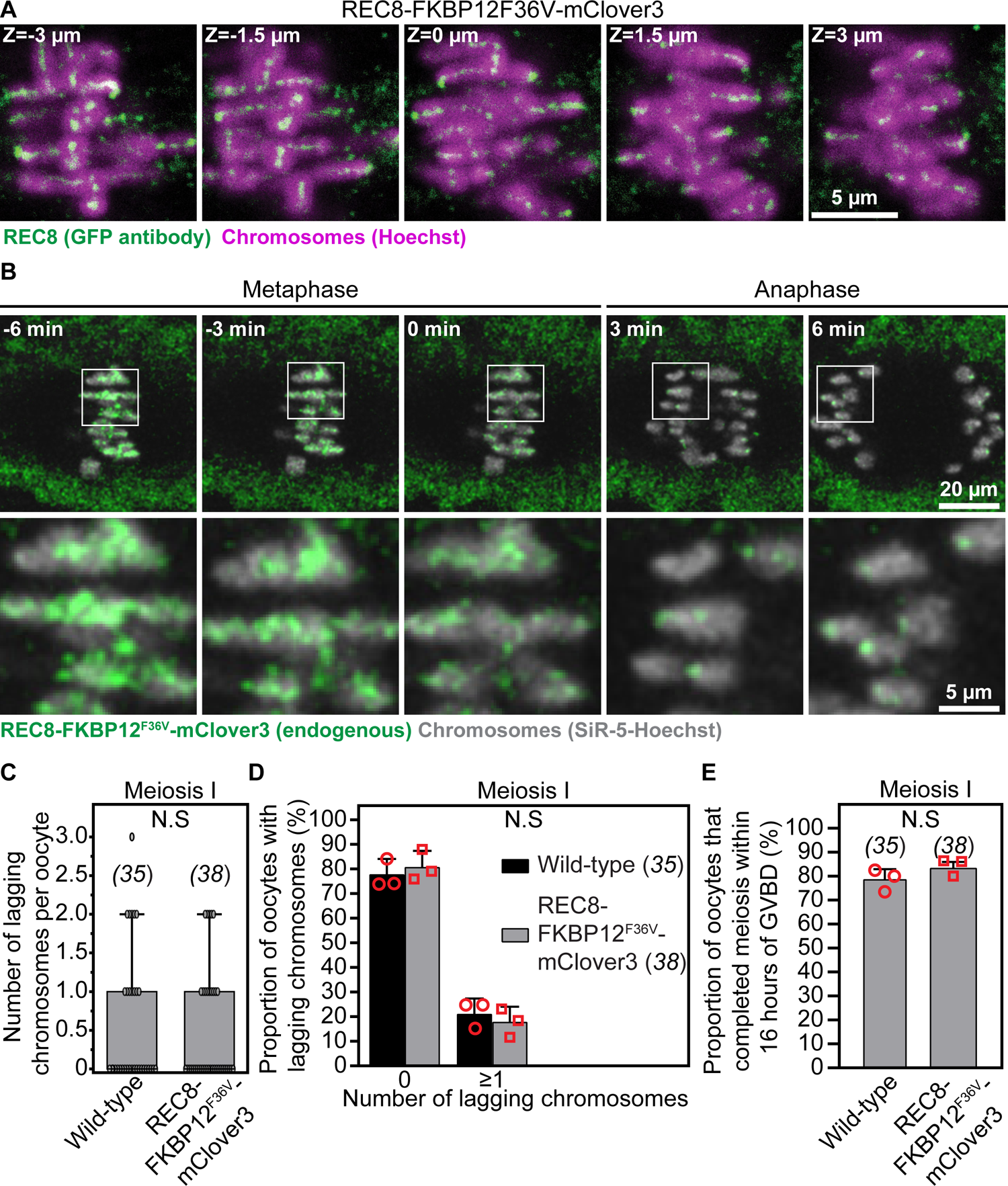
REC8-FKBP12^F36V^-mClover3 knockin mice enable high-resolution live imaging of endogenous REC8 dynamics during oocyte meiosis. (A) Representative confocal immunofluorescence microscopy images of REC8 (marked with GFP antibodies) and chromosomes (Hoechst) in metaphase I-stage oocytes of REC8-FKBP12^F36V^-mClover3 mouse. Single confocal sections spaced 1.5 µm apart are shown. (B) Images from a representative high-resolution time lapse movie of REC8-FKBP12^F36V^-mClover3 (endogenous REC8) and chromosomes (SiR-5-Hoechst) showing chromosome arm-specific removal of REC8 cohesin (and its retention at centromeric regions) during anaphase of meiosis I. (C) Quantification of the number of anaphase I lagging chromosomes per cell in wild-type and REC8-FKBP12^F36V^-mClover3 oocytes. Data points represent individual oocytes. (D) Quantification of the proportion of oocytes with 0 or more anaphase I lagging chromosomes in wild-type and REC8-FKBP12^F36V^-mClover3 oocytes. (E) Quantification of the proportion of oocytes that completed meiosis within 16 hours of GVBD in wild-type and REC8-FKBP12F36V-mClover3. Data are from 3 independent experiments. Statistical significance was evaluated using Student’s t-test (C-E). The number of analyzed oocytes is indicated in brackets and italics.

We further determined that endogenously tagged REC8 protein maintains its physiological function in chromosome cohesion using quantitative high-resolution live imaging of meiotic spindles and chromosomes (*36*) in oocytes isolated from homozygous knockin and wild-type mice. Analyses of chromosome organization and dynamics in these imaging assays showed that C-terminal tagging of endogenous REC8 with FKBP12^F36V^-mClover3 did not interfere with accurate metaphase I and II chromosome alignment, anaphase I chromosome segregation, or timing and efficiency of meiosis completion (Figs. 1, C-E and S2A; Movies S3 and S4). Cohesion dysfunction is associated with premature dissociation of bivalents into univalents (meiosis I) and of sister chromatids into single chromatids (meiosis II) (*17, 18, 20*), which is expected to increase the frequency of chromosome misalignment and missegregation. Our data thus demonstrate that REC8 cohesin function is not compromised in REC8-FKBP12^F36V^-mClover3 homozygous females.

To test whether endogenous REC8 cohesin in engineered animals can be targeted for rapid degradation via PROTACs, we acutely treated metaphase II-arrested eggs from reproductively young REC8-FKBP12^F36V^-mClover3 females with DMSO (Control) or dTAG-13 (*30, 31*). Using quantitative immunofluorescence microscopy of GFP antibody-stained cells to measure REC8 protein levels, we observed that REC8 cohesin fluorescence intensity was significantly reduced in dTAG-13 treated eggs (Fig. 2, A and B). Consistently, western blotting analyses showed robust degradation of REC8 protein in dTAG-13-treated cytosolic extracts of spermatocytes isolated from REC8-FKBP12^F36V^-mClover3 homozygous males (Fig. S3A).

**Fig. 2.**
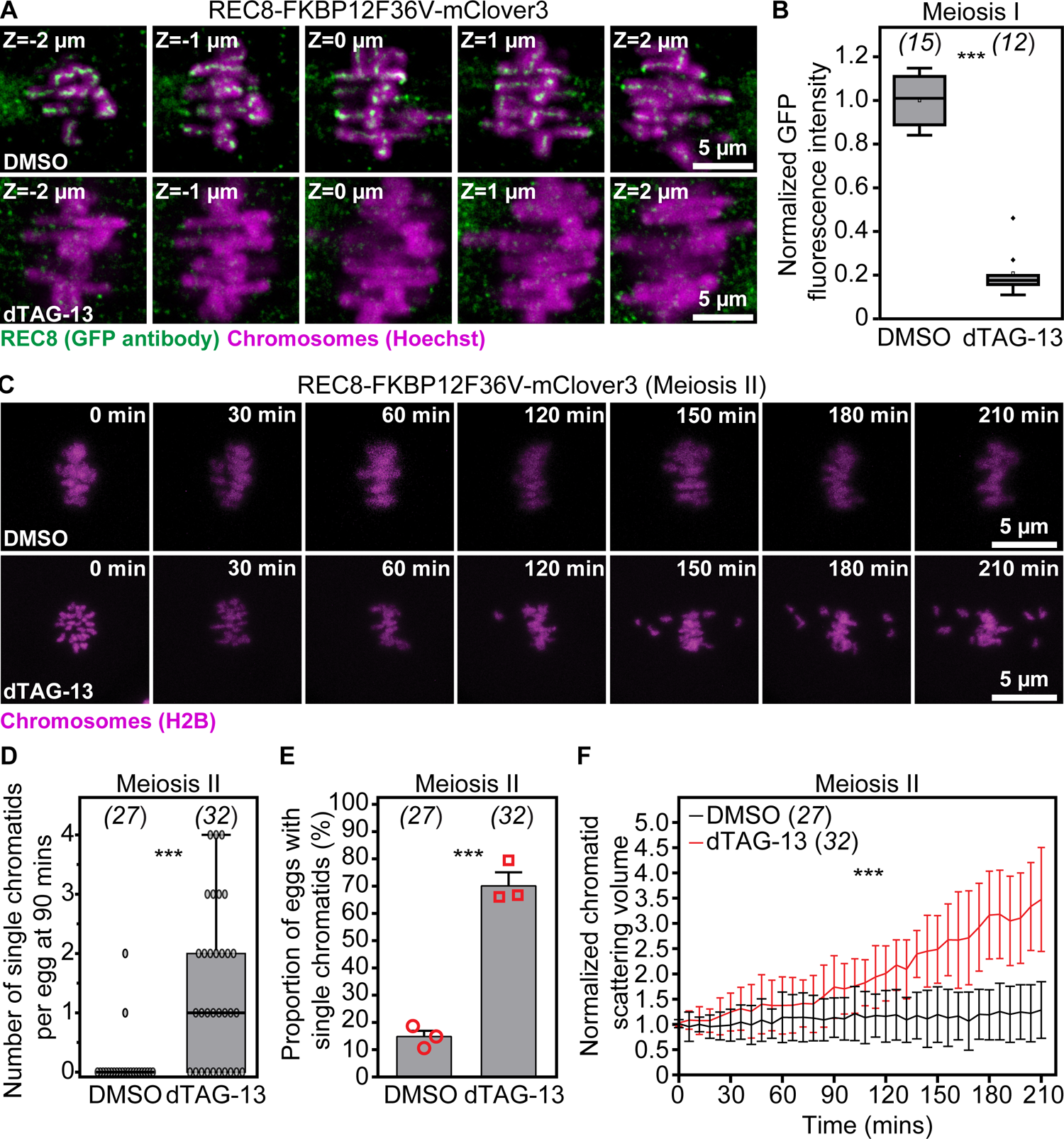
dTAG-13-mediated reduction of endogenous REC8 in REC8-FKBP12^F36V^-mClover3 oocytes rapidly recapitulates premature sister chromatid separation events associated with female reproductive aging. (A) Endogenous REC8 (labeled with GFP antibody) and metaphase I chromosomes (Hoechst) in DMSO- or dTAG-13-treated oocytes isolated from REC8-FKBP12^F36V^-mClover3 mice. Single confocal sections spaced 1 µm apart are shown. (B) Quantification of REC8 (labeled with GFP antibody) fluorescence intensity in DMSO- or dTAG-13-treated metaphase I oocytes isolated from REC8-FKBP12^F36V^-mClover3 mice. (C) Images from representative time-lapse movies of sister chromatids (H2B-mScarlet) in DMSO- or dTAG-13-treated eggs matured *in vitro* from REC8-FKBP12^F36V^-mClover3 oocytes. T = 0 min denotes the start of live imaging experiment. (D) Quantification of the number of single chromatids per egg at 90 mins of live imaging in DMSO- or dTAG-13-treated eggs matured *in vitro* from REC8-FKBP12^F36V^-mClover3 oocytes. (E) Quantification of the proportion of cells containing single chromatids in DMSO- or dTAG-13-treated eggs matured *in vitro* from REC8-FKBP12^F36V^-mClover3 oocytes. (F) Measurements of chromatid scattering volumes from time lapse movies of DMSO- or dTAG-13-treated metaphase II-arrested eggs matured *in vitro* from REC8-FKBP12^F36V^-mClover3 oocytes. Lines represent mean values and error bars indicate SEM. Data are from 3 independent experiments. Statistical significance was evaluated using Student’s t-test (B, D and E) or two-way ANOVA (F). The number of analyzed oocytes is indicated in brackets and italics.

We next examined whether PROTAC-mediated targeted degradation of REC8 cohesin can be used to induce aging-like meiotic chromatid separation events in reproductively young females. Here, we leveraged our established quantitative high-resolution live imaging assays of chromosome dynamics (*2, 3, 36*) to track in 3D the scattering of prematurely separated sister chromatids. We found that dTAG-13-mediated REC8 cohesin reduction could generate up to 4 prematurely separated single chromatids within 1.5 hours of treatment (Fig. 2, C-F; Movies S5 and S6), a process that normally takes years to decades in mice and humans (*2, 5, 20, 37–39*). Importantly, this rate is consistent with our previously reported measurements of premature chromatid separation in naturally aged eggs (*2*). Our quantitative analyses of chromatid separation additionally showed rapid scattering of sister chromatids after 1.5 hours of dTAG-13 treatment (Fig. 2F), which is consistent with sustained degradation of REC8 in the presence of dTAG-13. These results established that experimental reduction of REC8 in engineered REC8-FKBP12^F36V^-mClover3 mice rapidly generates aneuploidy-causing chromosomal defects that are common in eggs of reproductively older females.

The TRIM-Away method of antibody-based protein degradation (*2, 33, 40, 41*) can be coupled with GFP nanobodies to degrade GFP-tagged endogenous proteins (*33*). Our high-resolution live imaging data showed that GFP nanobodies also efficiently recognize exogenously expressed mClover3-tagged proteins in mouse oocytes (Fig. S4A). We thus predicted that GFP nanobody-coupled TRIM-Away of REC8 cohesin in REC8-FKBP12^F36V^-mClover3 mouse eggs could be used to induce aging-like premature separation of sister chromatids (Fig. 3A). To test this, we first microinjected exogenous TRIM21 expressing metaphase II stage eggs from engineered mice with mRNAs encoding GFP (*42*) or Gephyrin (*43*) (Control) nanobodies, then visualized chromatid dynamics at high spatiotemporal resolution. Analyses of chromosome dynamics in these experiments showed that GFP nanobody-mediated TRIM-Away of endogenous REC8 accelerated premature separation and scattering of sister chromatids (Fig. 3, B-E; Movies S7 and S8). In addition, we observed up to 4 prematurely separated chromatids 1.5 hours after REC8 degradation (Fig. 3C). These quantitative analyses confirmed that engineered REC8-FKBP12^F36V^-mClover3 mice offer multiple opportunities to rapidly generate aging-like aneuploidies in young eggs via chemical or genetic manipulation approaches.

**Fig. 3.**
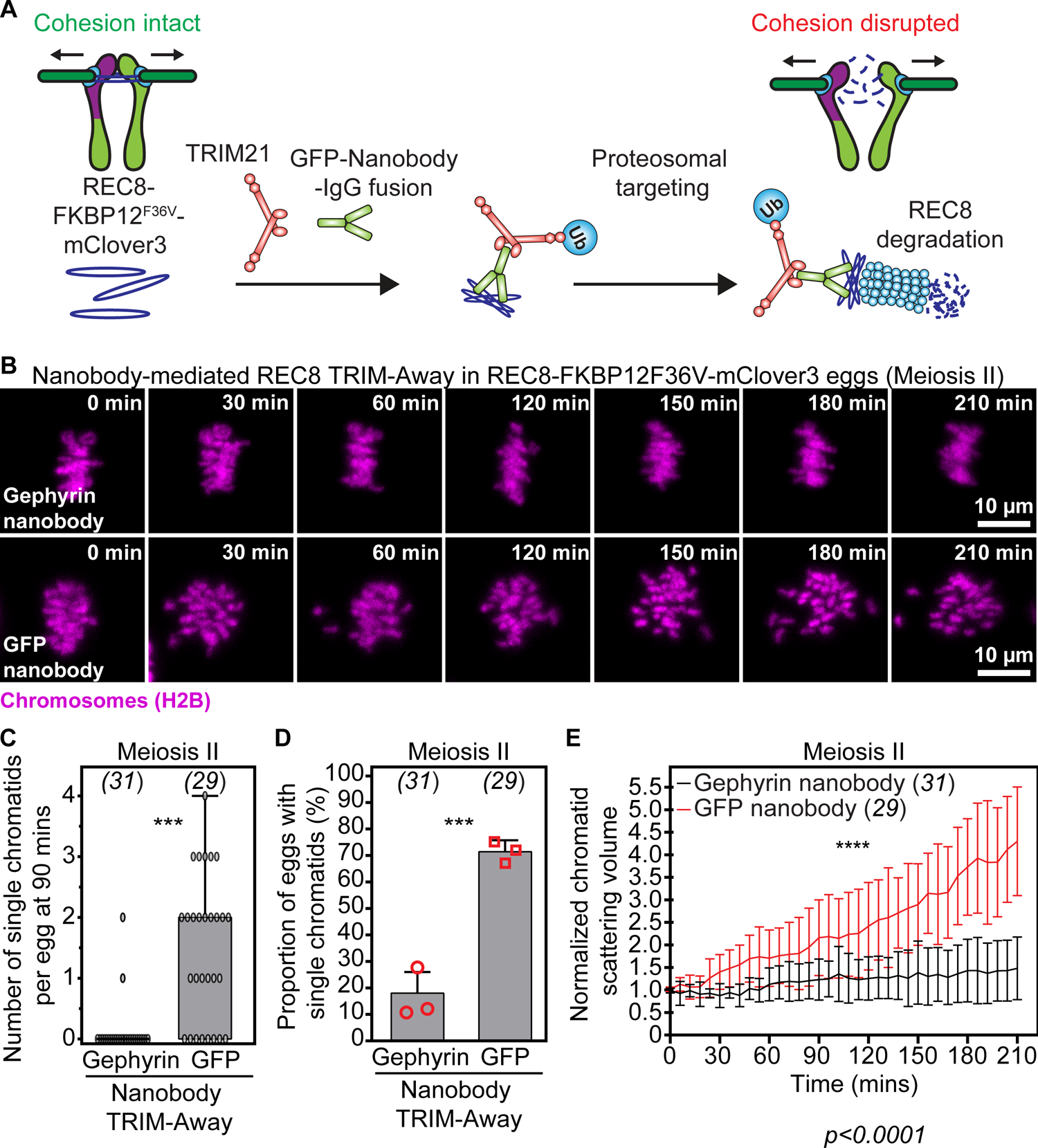
Nanobody-mediated TRIM-Away of endogenous REC8 in REC8-FKBP12^F36V^-mClover3 oocytes accelerates premature separation of sister chromatids. (A) Graphical description of nanobody-based approach for rapid degradation of endogenous REC8 in REC8-FKBP12^F36V^-mClover3 oocytes. Recognition of mClover3-tagged REC8 by IgG-Fc1 fusion GFP nanobodies in TRIM21 expressing REC8-FKBP12^F36V^-mClover3 eggs induces endogenous REC8 degradation via TRIM-Away and generates prematurely separated sister chromatids. (B) Images from representative time-lapse movies of sister chromatids (H2B-mScarlet) in Gephyrin or GFP TRIM-Away eggs from REC8-FKBP12^F36V^-mClover3 mice. T = 0 min denotes the start of live imaging experiment. (C) Quantification of the number of single chromatids per egg at 90 mins of live imaging in Gephyrin or GFP TRIM-Away eggs from REC8-FKBP12^F36V^-mClover3 mice. (D) Quantification of the proportion of cells containing single chromatids in Gephyrin or GFP TRIM-Away eggs from REC8-FKBP12^F36V^-mClover3 mice. (E) Measurements of chromatid scattering volumes from time lapse movies of Gephyrin or GFP TRIM-Away metaphase II-arrested eggs from REC8-FKBP12^F36V^-mClover3 mice. Lines represent mean values and error bars indicate SEM. Data are from 3 independent experiments. Statistical significance was evaluated using Student’s t-test (C and D) or two-way ANOVA (E). The number of analyzed oocytes is indicated in brackets and italics.

We previously identified that female reproductive aging is accompanied by meiotic spindle-specific disruption of the actin cytoskeleton, which predisposes mammalian eggs to premature chromatid separation (*2*). To confirm that our new experimental system of female reproductive age-related aneuploidy recapitulates this new cytoskeletal function, we first treated metaphase II-arrested eggs with dTAG-13 to reduce chromosome cohesion and then disrupted F-actin with Cytochalasin D (*2, 36*). Consistent with our previous finding (*2*), quantification of chromatid scattering volume in our high spatiotemporal resolution 3D live imaging assays of chromosome dynamics (*2, 36*) revealed that F-actin disruption alone caused gradual splitting of sister chromatids in reproductively young REC8-FKBP12^F36V^-mClover3 females (Fig. 4, A-D; Movies S9 and S10), with an average of 1 single chromatid evident 1.5 hours after drug addition (Fig. 4B). Confirming our initial data, dTAG-13-mediated cohesion disruption caused significant chromatid separation and scattering (Fig. 4, A-C; Movies S11), with up to 3 single chromatids present on the spindle after 1.5 hours of PROTAC treatment (Fig. 4B). Validating that this experimental system indeed recapitulates aneuploidies associated with female reproductive age-related F-actin dysfunction, Cytochalasin D addition accelerated the scattering of prematurely separated sister chromatids in dTAG-13 treated eggs containing reduced cohesion (Fig. 4D; Movie S12).

**Fig. 4.**
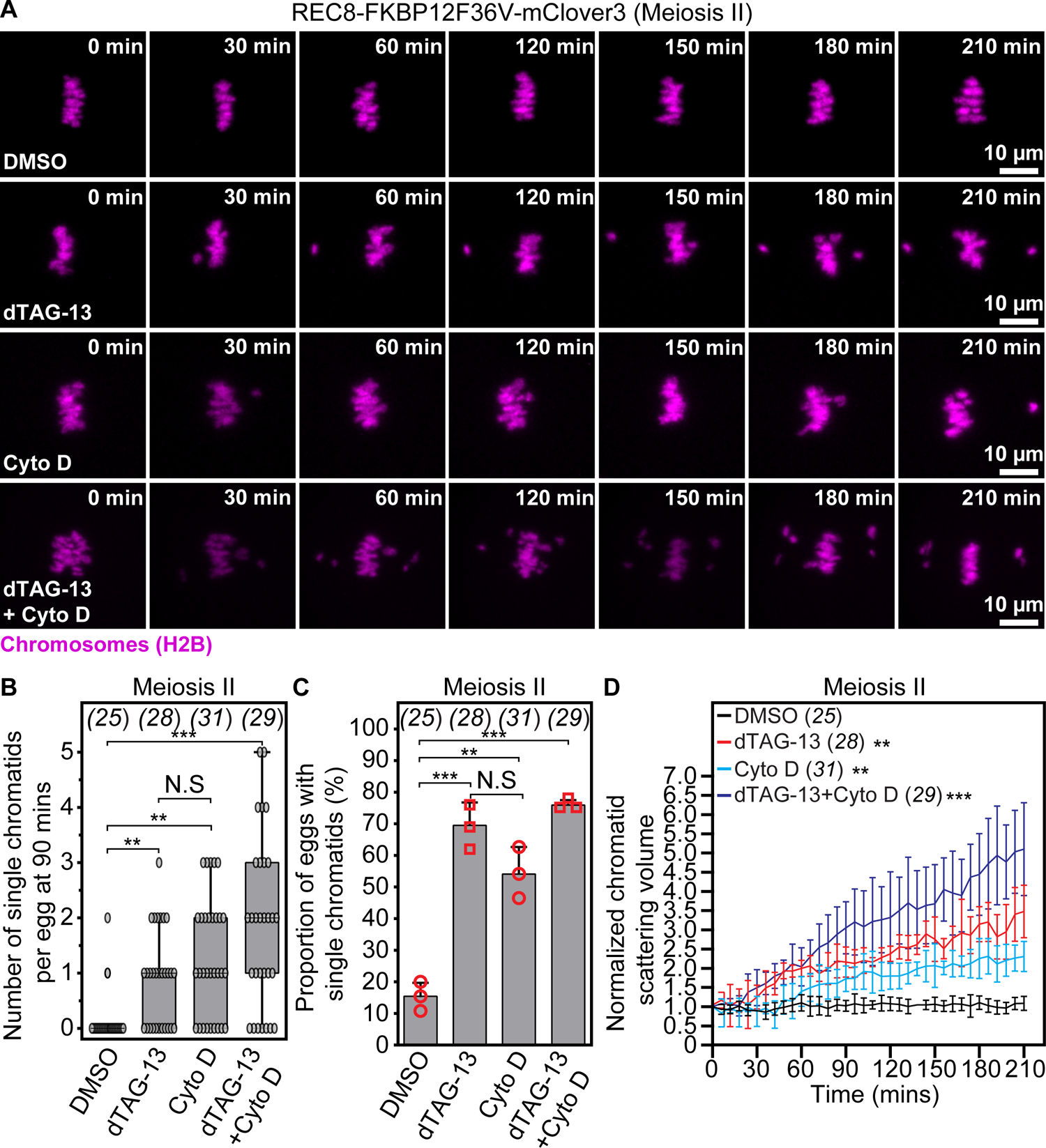
Premature sister chromatid separation arising from PROTAC-mediated REC8 cohesin degradation is accelerated by actin cytoskeleton disruption. (A) Images from representative time-lapse movies of sister chromatids (H2B-mScarlet) in DMSO-, dTAG-13-, Cytochalasin D (CytoD)- or dTAG-13 and CytoD-treated eggs matured *in vitro* from REC8-FKBP12^F36V^-mClover3 oocytes. T = 0 min denotes the start of live imaging experiment. (B) Quantification of the number of single chromatids per egg at 90 mins of live imaging in DMSO-, dTAG-13-, CytoD- or dTAG-13 and CytoD-treated eggs matured *in vitro* from REC8-FKBP12^F36V^-mClover3 oocytes. (C) Quantification of the proportion of cells containing single chromatids in DMSO-, dTAG-13-, CytoD- or dTAG-13 and CytoD-treated eggs matured *in vitro* from REC8-FKBP12^F36V^-mClover3 oocytes. (D) Measurements of chromatid scattering volumes from time lapse movies of DMSO-, dTAG-13-, CytoD- or dTAG-13 and CytoD-treated eggs matured *in vitro* from REC8-FKBP12^F36V^-mClover3 oocytes. Lines represent mean values and error bars indicate SEM. Data are from 3 independent experiments. Statistical significance was assessed using Student’s t-test (B and C) or two-way ANOVA (D). The number of analyzed oocytes is provided in brackets and italics.

A small molecule-based tool to directly manipulate chromosome cohesion holds strong promise for rapid screening and discovery of proteins whose age-related disruption can predispose eggs to aneuploidy, independently of cohesion loss. We thus tested whether our PROTAC-based experimental system of oocyte aging can be applied to discover factors associated with the remarkable rise in egg aneuploidy at advanced reproductive ages (*4–6*). The histone H3 variant CENP-A (*44*) is required for proper centromere function in oocytes, fertility and transmission of the genome to the next generation (*45, 46*). Reproductive age-related depletion of CENP-A and other centromeric components was recently proposed to underlie kinetochore dysfunction and chromosome segregation errors in aged oocytes (*47*). However, it is unknown whether CENP-A loss can trigger the sharp increase in the incidence of egg aneuploidy observed during maternal aging (*4–6*). We addressed this by first treating exogenous TRIM21 expressing REC8-FKBP12^F36V^-mClover3 metaphase II stage eggs for 3 hours with DMSO (Control) or dTAG-13. We then performed Control or CENP-A TRIM-Away by microinjecting mouse IgG or CENP-A heavy chain antibodies into DMSO- or dTAG-13-treated eggs. Immunofluorescence microscopy of metaphase chromosome spreads showed that CENP-A protein abundance was successfully reduced by approximately 30% after TRIM-Away (Fig. 5, B and C), a value comparable to CENP-A depletion levels in naturally aged eggs (*47*). dTAG-13 treatment of Control IgG TRIM-Away eggs induced modest separation and scattering of sister chromatids at 2 hours of treatment (Figs. 5, A and D, S5A; Movies S13 and S14). TRIM-Away-mediated aging-like degradation of CENP-A in DMSO-treated control eggs led to chromatid scattering events comparable to dTAG-13-mediated partial removal of cohesion (Figs. 5, A and D, S5A; Movie S15). Since CENP-A is critical for the identity and function of centromeres that also serve as sole regions of meiosis II chromosome cohesion (*5, 17, 18, 20, 48*), failure to maintain centromeric cohesion in CENP-A TRIM-Away eggs could explain this observation. In stark contrast, CENP-A TRIM-Away in dTAG-13-treated eggs containing reduced cohesion generated an average of 3-4 prematurely separated chromatids and accelerated widespread scattering of chromosomes on the spindle (Figs. 5, A and D, S5A; Movie S16). These results indicate that maternal age-related depletion of CENP-A and cohesion proteins serve as independent sources of oocyte aneuploidies associated with female reproductive aging. In identifying CENP-A as a protein whose dysfunction likely causes exponential rise in age-related egg aneuploidy, these data also demonstrated that our PROTAC-based experimental system can be implemented to rapidly identify key factors that impact female reproductive longevity.

**Fig. 5.**
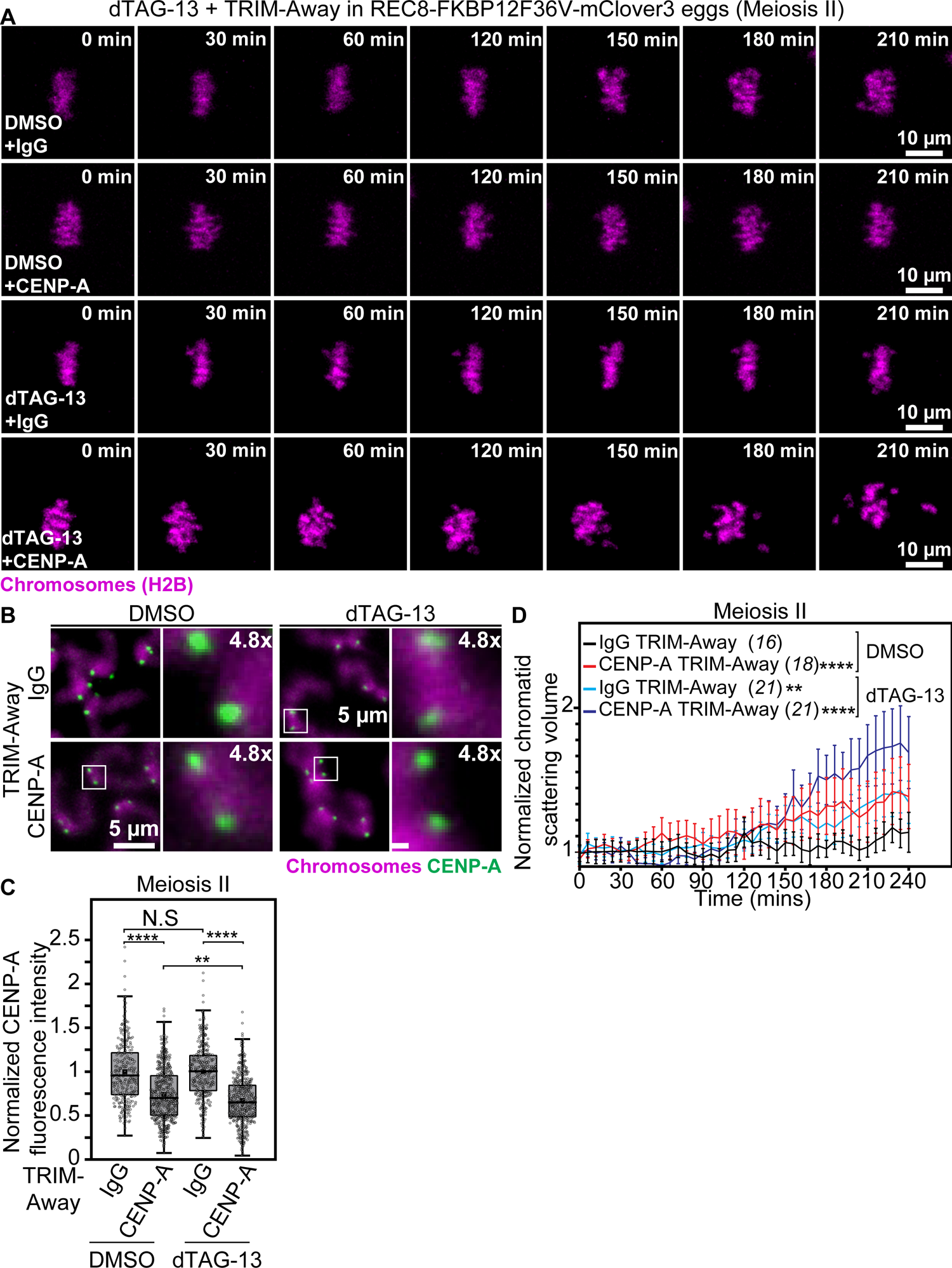
Aging-like CENP-A depletion triggers premature separation of sister chromatids independently of cohesion loss. (A) Images from representative time-lapse movies of sister chromatids (H2B-mScarlet) in DMSO-treated and IgG TRIM-Away, DMSO-treated and CENP-A TRIM-Away, dTAG-13-treated and IgG TRIM-Away or dTAG-13-treated and CENP-A TRIM-Away eggs from REC8-FKBP12^F36V^-mClover3 mice. T = 0 min denotes the start of live imaging experiment. (B) Representative maximum intensity projected immunofluorescence images of CENP-A and chromosomes in metaphase II chromosomal spreads of DMSO-treated and IgG TRIM-Away, DMSO-treated and CENP-A TRIM-Away, dTAG-13-treated and IgG TRIM-Away or dTAG-13-treated and CENP-A TRIM-Away eggs from REC8-FKBP12^F36V^-mClover3 mice. Boxes mark regions of interest that are 4.8x magnified. (C) Quantification of CENP-A fluorescence intensities in metaphase II chromosomal spreads of DMSO-treated and IgG TRIM-Away (295 centromeres), DMSO-treated and CENP-A TRIM-Away (535 centromeres), dTAG-13-treated and IgG TRIM-Away (347 centromeres) and dTAG-13-treated and CENP-A TRIM-Away (465 centromeres) eggs from REC8-FKBP12^F36V^-mClover3 mice. (D) Measurements of chromatid scattering volumes from time lapse movies of DMSO-treated and IgG TRIM-Away, DMSO-treated and CENP-A TRIM-Away, dTAG-13-treated and IgG TRIM-Away or dTAG-13-treated and CENP-A TRIM-Away eggs from REC8-FKBP12^F36V^-mClover3 mice. Lines represent mean values and error bars indicate SEM. Data are from three independent experiments. Statistical significance was assessed using two way ANOVA (C and D). The number of analyzed oocytes is provided in brackets and italics.

In this study, we have developed a versatile experimental system for induction of aging-like chromosomal phenotypes in young eggs. By overcoming the need to age mice or purchase aged animals, this drug-based cohesion manipulation system provides a fast and cost-effective approach to discover the causes of reproductive age-related egg aneuploidy. We anticipate that innovative live imaging assays of REC8 protein dynamics enabled by this experimental system will also provide new opportunities to probe the mechanisms of chromosome cohesion in female meiosis. Next steps will include combining this approach with libraries of small molecules, PROTACs and clinical compounds for unbiased identification and mechanistic studies of new meiotic fidelity proteins. In future, clinical manipulation of meiotic processes uncovered via this approach could accelerate next-generation fertility treatments that reduce the incidence of aneuploidy in fertilizable eggs.

## Materials and Methods

### Mouse strains and husbandry

All protocols involving mice were approved by the Yale University Institutional Animal Care and Use Committee (approval number 2021-20408). Mice were housed under specific pathogen-free (SPF) conditions with a 12-hour light/dark cycle. The mouse strains used in this study included REC8-FKBP12^F36V^-mClover3 and wild-type C57BL/6J (The Jackson Laboratory).

### CRISPR-Cas generation of REC8-FKBP12^F36V^-mClover3 mice

The Rec8 FKBP12-F36V-mClover3 mouse model was generated via CRISPR/Cas9-mediated genome editing (*49–51*). Potential Cas9 target guide (protospacer) sequences near the REC8 translation termination site were screened using the online tool CRISPOR http://crispor.tefor.net (*52*) and candidates were selected. crRNAs containing the chosen sequences were synthesized (IDT). crRNA/tracrRNA/Cas9 RNPs were complexed and tested for activity by zygote electroporation, incubation of embryos to blastocyst stage, and genotype scoring of indel creation at the target sites. A gRNA that demonstrated high activity and that directed Cas9 cleavage to the translation termination codon, 5’ GGTTTAATGAACTTGCTCAG, was selected for creating the knockin allele. Accordingly, a 1.5kb long single-stranded DNA (lssDNA) recombination template containing the FKBP12^F36V^-mClover3 element was synthesized (IDT). The injection mix of gRNA/Cas9 RNP + lssDNA was microinjected into the pronuclei of C57Bl/6J zygotes (*51*). Embryos were transferred to the oviducts of pseudo pregnant CD-1 foster females using standard techniques (*53*). Genotype screening of tissue biopsies from founder pups was performed by PCR amplification and Sanger sequencing to verify the knockin allele. Germline transmission of the correctly targeted allele was confirmed by breeding and sequence analysis. To maintain genetic robustness, the REC8-FKBP12^F36V^-mClover3 colony was outbred to C57BL/6J mice every five generations. Following the establishment of homozygosity, this strain was maintained as an independent line for subsequent experiments.

### Genotyping of REC8-FKBP12^F36V^-mClover3 mice

Genotyping was performed using DNA extracted from ear punch biopsies. DNA was amplified using the KAPA HiFi HotStart ReadyMix (Roche, 07958935001) according to the manufacturer’s instructions. PCR primers specific for REC8 (see Table S1, BM729-BM735) were used to distinguish between alleles. The expected product sizes were 550 bp for the wild-type (WT) allele and 1700 bp for the REC8-FKBP12^F36V^-mClover3 allele.

### Mouse oocyte isolation, maturation, culture, and microinjection

Oocytes were collected from ovaries of 6–8-week-old REC8-FKBP12^F36V^-mClover3 female mice. Isolated oocytes were matured *in vitro* and microinjected with 6–8 picoliters of *in vitro* transcribed mRNA as previously described (*36*).

### Generation of *in vitro* mRNA transcription constructs and mRNA synthesis

Oligonucleotide sequences used for plasmid construction are provided in Table S1. All synthesized gene sequences were obtained from Twist Bioscience. To generate pGEM-H2B-mScarlet, the coding sequence of mouse H2B (obtained by gene synthesis) was amplified with KS109 and KS110 and transferred into pGEM-mScarlet backbone that was linearized with EcoRI and XhoI. pGEM-MAP4-MTBD-mClover was generated by Gibson assembly (New England Biolabs) of KS111 and KS112 amplified synthetic mclover3 sequence into the AflII and NcoI site of pGEM-EGFP-MAP4-MTBD (*2*). pGEM-mNeonGreen-MAP4-MTBD was constructed by Gibson assembly of KS158 and KS159 amplified synthetic mNeonGreen sequence into pGEM-MAP4-MTBD-mClover3 that was linearized with KS156 and KS157. Generation of pGEM-SNAP-TRIM21 construct was described previously (*2*). pGEM-VHHGFP4-hIgG1-Fc1 plasmid was constructed using Gibson assembly. A PCR fragment amplified with primers BM343 and BM344 from the pcDNA3-Nslmb-VHHGFP4 (*42*) (Addgene Plasmid #35579) plasmid was inserted into the HindIII–EcoRI site of the pGEM-hIgG1-Fc1 vector (*3*). pGEM-VHHGFP4-mScarlet was generated by linearizing pGEM-HE(*54*) with BM715 and BM811, and transferring into it using Gibson assembly BM812 and BM813 flanked synthesized coding sequence of VHHGFP4. Gephyrin nanobody sequence (*43*) was obtained by gene synthesis. pGEM-GephyrinVHH-hlgG1-Fc1 was generated by transferring JL7 and JL8 amplified coding sequence of Gephyrin nanobody into the HindIII–EcoRI site of pGEM-hIgG1-Fc1. pGEM-GephyrinVHH-mScarlet was generated by linearizing pGEM-HE with JL9 and JL10 and transferring into it by Gibson assembly JL3 and JL4 flanked synthetic coding sequence of Gephyrin VHH. pGEM-mScarlet-NLS was constructed by amplifying the coding sequence of mScarlet-NLS (synthesized DNA) using primers BM432 and BM433 and transferring it into the HindIII site of pGEM-HE using Gibson assembly. pGEM-mClover3-NLS was generated using Gibson assembly to transfer BM432 and FG40 flanked mClover3 coding sequence into pGEM-mScarlet-NLS that was linearized with FG39 and BM494. Capped mRNAs were synthesized using a T7 polymerase (mMessage mMachine kit, Ambion). mRNA concentrations were measured using a NanoDrop spectrophotometer (Thermo Fisher Scientific).

### Drug addition and fluorescence labeling experiments

Stock solutions of all small-molecule inhibitors were prepared in dimethyl sulfoxide (DMSO). To disrupt actin, oocytes were treated with Cytochalasin D (C8273-1MG, Merck) at a final concentration of 5 μg/ml in M2 medium for 1 hour. For rapid protein degradation experiments, eggs were exposed to 3 μM dTAG-13 (SML2601, Merck). All compounds were dissolved in DMSO (D2650-5X5ML, Merck) and diluted in M2 medium. In live imaging experiments, chromosomes were visualized by treating cells for 1 hour with 100 nM SiR-5-Hoechst dye diluted in M2 medium. DMSO was used as control samples at a final concentration of 0.003% in M2 medium.

### Western blotting

Protein samples were prepared by washing sperm in PBS and lysing in LDS sample buffer (NuPAGE; Invitrogen). Samples were agitated thoroughly and snap-frozen at −80°C until use. Before electrophoresis, samples were boiled at 95°C for 10 minutes with reducing agent (NuPAGE). Proteins were separated on 4–12% Bis-Tris gels (NuPAGE; Invitrogen) and transferred to PVDF membranes (Immobilon-P; Millipore). Membranes were blocked with 3% BSA in PBS for 1 hour at room temperature. Primary antibody incubation was performed overnight at 4°C in 5% (w/v) milk-PBST (PBS containing 0.05% (w/v) Tween 20) blocking buffer with GFP antibody (1:1000, Novus Bio, NBP2-50059). Membranes were washed for 5 minutes three times with PBST then incubated with a goat anti-rabbit IgG StarBright Blue 700 secondary antibody (1:5000, Bio-Rad) prepared in blocking buffer for 1 hour at room temperature. Actin-Rhodamine (Bio-Rad) was used as a loading control to detect actin. Following three additional washes in PBST, fluorescence signals were detected using the ChemiDoc MP Imaging System (Bio-Rad) with auto-exposure mode.

### Fixation and Immunostaining of Mouse Oocytes and Eggs

Oocytes were fixed in a solution containing 100 mM Hepes, 10 mM MgSO4, 50 mM EGTA, 0.5% Triton X-100 (v/v), and 2% formaldehyde (v/v) at 37°C for 30 minutes. Fixed cells were blocked overnight at 4°C in PBT (PBS containing 0.3% Triton X-100 (v/v)) and 3% BSA (w/v). Primary antibody (GFP, 1:100; Roche, 11814460001) was applied overnight at 4°C. Cells were washed and incubated with Alexa Fluor 488–labeled anti-mouse secondary antibody (1:500; Molecular Probes) and Hoechst 33342 (5 μg/ml; Molecular Probes) for 1 hour at room temperature.

### Metaphase chromosomal spreading, fixation, and immunostaining

Metaphase II-arrested eggs were prepared for chromosomal spreading by removing the zona pellucida using Tyrode’s acid solution (Sigma, T1788). Eggs were then thoroughly washed through droplets of fresh M2 medium and recovered by incubating for at least 10 minutes at 37°C. For chromosomal spreading, single cells were placed on a 15-well multitest slide (MP Biomedical, 096041505) in a fixative solution containing 1% formaldehyde (v/v), 0.15% Triton X-100 (v/v), and 3% dithiothreitol (v/v), pH 9.2–9.4. Slides were incubated overnight in a humidified chamber at room temperature. The next day, chromosomal spreads were air-dried and blocked in PBS containing 3% BSA (w/v) for 30 minutes. Spreads were then incubated with λ-phosphatase (New England Biolabs, P0753S) at 30°C for 2 hours. After washing in PBS, immunostaining was performed using CENP-A primary antibody (Cell Signaling, C51A7) at 37°C for 1.5 hours, followed by Alexa Fluor 488–labeled anti-rabbit secondary antibody (1:500; Invitrogen, A11008) for 1 hour at 37°C and Hoechst 33342 (5 μg/ml; Thermo Scientific, 62249) for 30 minutes at room temperature. Slides were washed and mounted with VECTASHIELD Antifade Mounting Medium (2B Scientific, H-1000-10) under 22 × 22 mm glass coverslips (Epredia, 22X22-1-002G), which were sealed with nail varnish.

### TRIM-Away-mediated protein degradation in dTAG-13-treated mouse eggs

For nanobody-mediated TRIM-Away of Gephyrin or GFP proteins in dTAG-13-treated eggs, metaphase II–arrested eggs expressing TRIM21 were microinjected in M2 medium with 2–3 picoliters of GFP or Gephyrin nanobody-encoding mRNAs.

In CENP-A degradation TRIM-Away experiments, TRIM21-expressing metaphase II-arrested eggs were incubated in 3 μM DMSO or dTAG13 for 1.5h before microinjection of control Rabbit-IgG (R&D Systems, A-105-C) or CENP-A antibodies (Cell Signaling, C51A7). Successful microinjection was confirmed by co-microinjection of fluorescent Dextran Texas Red (1:100 dilution; Invitrogen, D1829) in 0.05% (v/v) NP-40-PBS. All TRIM-Away microinjections and successive live imaging experiments were performed in M2 medium containing DMSO or dTAG

### High-resolution live cell imaging

Confocal time-lapse images of mouse oocyte and metaphase II–arrested eggs were acquired using a Zeiss LSM 800 Airyscan or LSM 900 Airyscan 2 microscope equipped with a 40× C-Apochromat 1.2 numerical aperture (NA) water-immersion objective and an environmental chamber maintained at 37°C. Image acquisition was controlled with ZEN2 software (Zeiss) and conducted at a temporal resolution of 6 minutes. REC8-mClover3 was imaged on a Leica STELLARIS 5 confocal laser scanning microscope (Leica) equipped with an environmental incubator box and a 40× C-Apochromat 1.2N.A. water-immersion objective and conducted at a temporal resolution of 3 minutes. Z-stacks were captured with a thickness of approximately 40 μm, with confocal sections spaced 1.5 μm apart. Oocytes were imaged in M2 medium under mineral oil as described previously (*36*).

### Immunofluorescence microscopy

Confocal immunofluorescence imaging was performed using a Zeiss LSM 800 or LSM 900 confocal microscope with a 40× C-Apochromat 1.2-NA water-immersion objective. Z-stacks were acquired with a thickness of 2.0 μm, with 0.5-μm confocal sections.

For GFP immunofluorescence imaging, images were were acquired in 1 μm steps over a 4 μm range at the midplane of meiotic chromosomes using the Airyscan module on the Zeiss LSM 800 or Airyscan 2 module LSM 900 microscopes. Post-acquisition super-resolution images were generated through 3D Airyscan processing using ZEN2 software (Zeiss). Eggs were imaged in M2 medium under mineral oil as described previously (*36*).

### Measurement of fluorescence intensity in metaphase chromosome spreading assays

Mean fluorescence intensity of REC8 on metaphase I chromosomes was measured in ImageJ by manually tracing regions of interest around maximum intensity projected images of chromosome spreads. Mean fluorescence intensity of CENP-A on metaphase II sister chromatids was measured in ImageJ by manually tracing centromeric regions of DNA in maximum intensity projected images of chromosome spreads.

### 3D surface reconstruction and measurement of chromatid scattering volume

Chromatid surfaces were reconstructed in 3D using Imaris (Bitplane) from high-resolution time lapse movies of H2B-mScarlet. Chromatid scattering volume was quantified using object-oriented bounding box analysis in Imaris, identifying the minimal cuboid volume enclosing all chromatids at each time point. For each egg, the scattering volume was normalized by dividing the bounding box volume at each time point by the volume at the start of the live imaging experiment (t = 0 min). This normalization enabled direct comparison of chromatid dynamics across samples.

### Statistical data analysis

All statistical analyses were conducted using GraphPad Prism software (GraphPad) or OriginPro software (OriginLab). Statistical box plots represent median (line), mean (small square), 5th, 95th (whiskers) and 25th and 75th percentile (box enclosing 50% of the data). Normalizations were performed by dividing individual data points in control and experimental groups by the average value of control group data points. Differences between two groups were analyzed using Student’s t-test, while comparisons among more than two groups were assessed using Two-way analysis of variance (ANOVA). Statistical significance is denoted as *P < 0.05, **P < 0.005, and ***P < 0.0005. Non-significant values are indicated as N.S.

## Data availability

All data are available in the main text or the supplementary materials. Plasmid constructs used in this study will be shared upon request and will be made publicly available through Addgene. REC8-FKBP12F36V-mClover3 mice are in the process of being deposited to Jackson Laboratories for sharing with the wider research community.

## Acknowledgments

We are grateful to Grazvydas Lukinavicius (Max Planck Institute for Multidisciplinary Sciences) for sharing 5-SiR-Hoechst dyes for labelling oocyte chromatin. We thank members of the Mogessie lab for critical discussions of this work. For the purpose of open access, the corresponding author has applied a CC BY public copyright license to any Author Accepted Manuscript version arising from this submission.

## Author contributions

Conceptualization: BM

Methodology: JL, BM

Investigation: JL, BM, TL, AK

Visualization: JL, BM, TL, AK, LT, XX, SB, TN

Funding acquisition: JL, BM

Project administration: BM

Supervision: BM

Writing – original draft: BM

Writing – review & editing: JL, BM

## Materials & Correspondence

Requests for materials should be addressed to

## Extended data

**Fig. S1.**
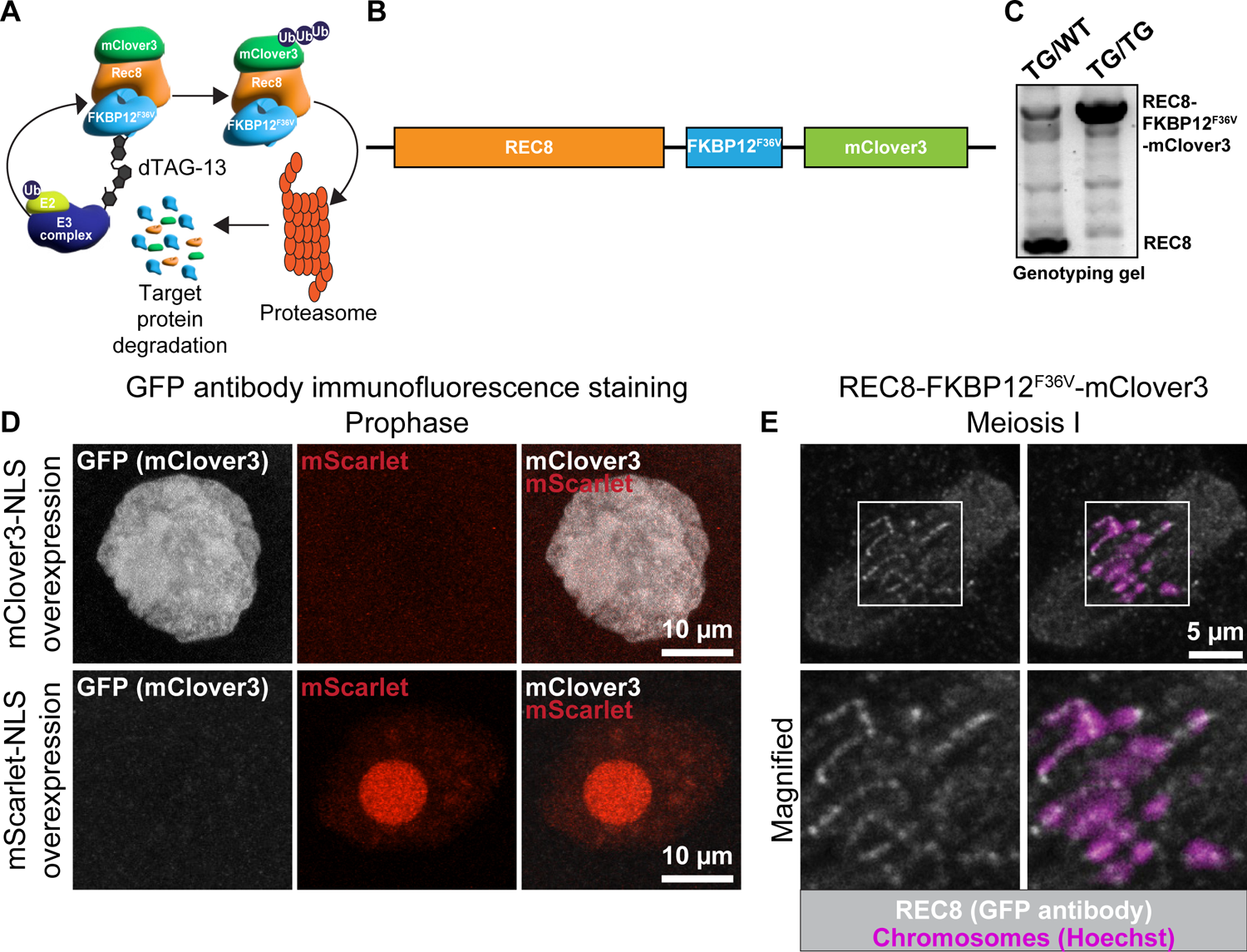
Generation of REC8-FKBP12^F36V^-mClover3 mice as a versatile cohesion manipulation system. (A) Graphical description of the principle of dTAG-13 proteolysis targeting chimera-mediated endogenous protein degradation. (B) Graphical description of gene arrangements in REC8-FKBP12^F36V^-mClover3 knockin mice. Boxes depicting genes are drawn to scale. (C) A representative genotyping agarose gel analysis of REC8-FKBP12^F36V^-mClover3 mouse genomic DNA using oligonucleotide pairs designed to distinguish between wild-type and knockin REC8 alleles. (D) Representative maximum intensity projected immunofluorescence images showing that GFP antibodies specifically recognize mClover3 but not mScarlet in mClover3-NLS or mScarlet-NLS expressing prophase-arrested wild-type oocytes. (E) Representative maximum intensity projected immunofluorescence images of REC8 (detected with GFP antibody) and chromosomes in metaphase I-stage oocytes isolated from REC8-FKBP12^F36V^-mClover3 mice. Boxes represent regions of interest magnified in lower panels.

**Fig. S2.**
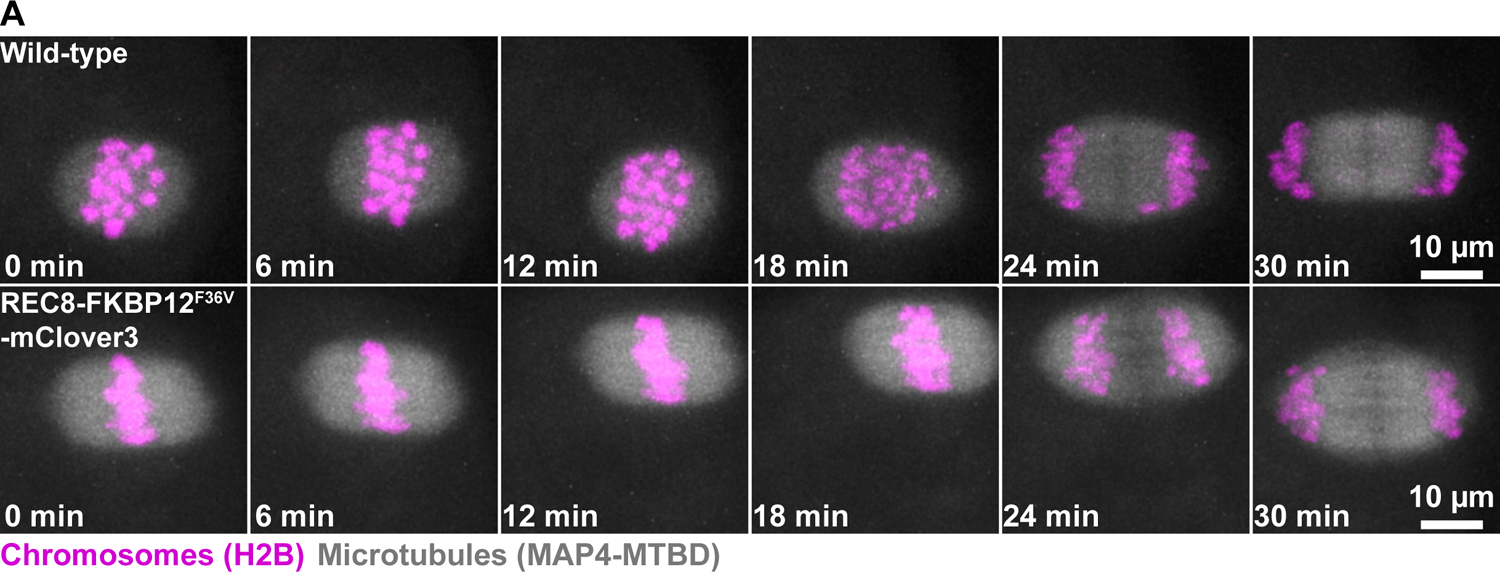
CRISPR-Cas based C-terminal tagging of endogenous REC8 does not disrupt accurate oocyte chromosome alignment and segregation. (A) Images from representative time lapse movies of chromosomes (marked with H2B-mScarlet) and meiotic spindles (marked with MAP4 microtubule-binding domain (mNeonGreen-MAP4-MTBD)) in wild-type and REC8-FKBP12^F36V^-mClover3 oocytes.

**Fig. S3.**
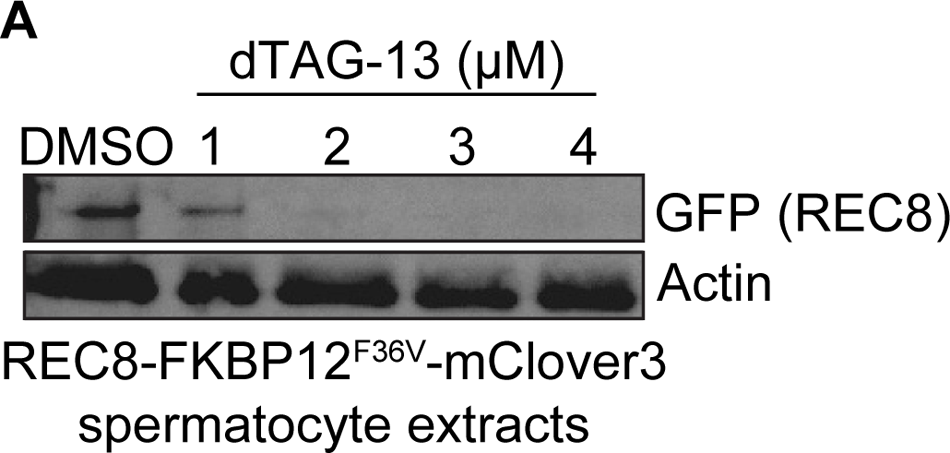
Validation of dTAG-13-mediated REC8 degradation in REC8-FKBP12^F36V^-mClover3 spermatocytes. (A) Western blotting analyses of REC8-FKBP12^F36V^-mClover3 spermatocyte extracts treated with DMSO (control) or varying concentrations of dTAG-13. GFP antibody was used for detection of REC8 protein. Actin was used as loading control.

**Fig. S4.**
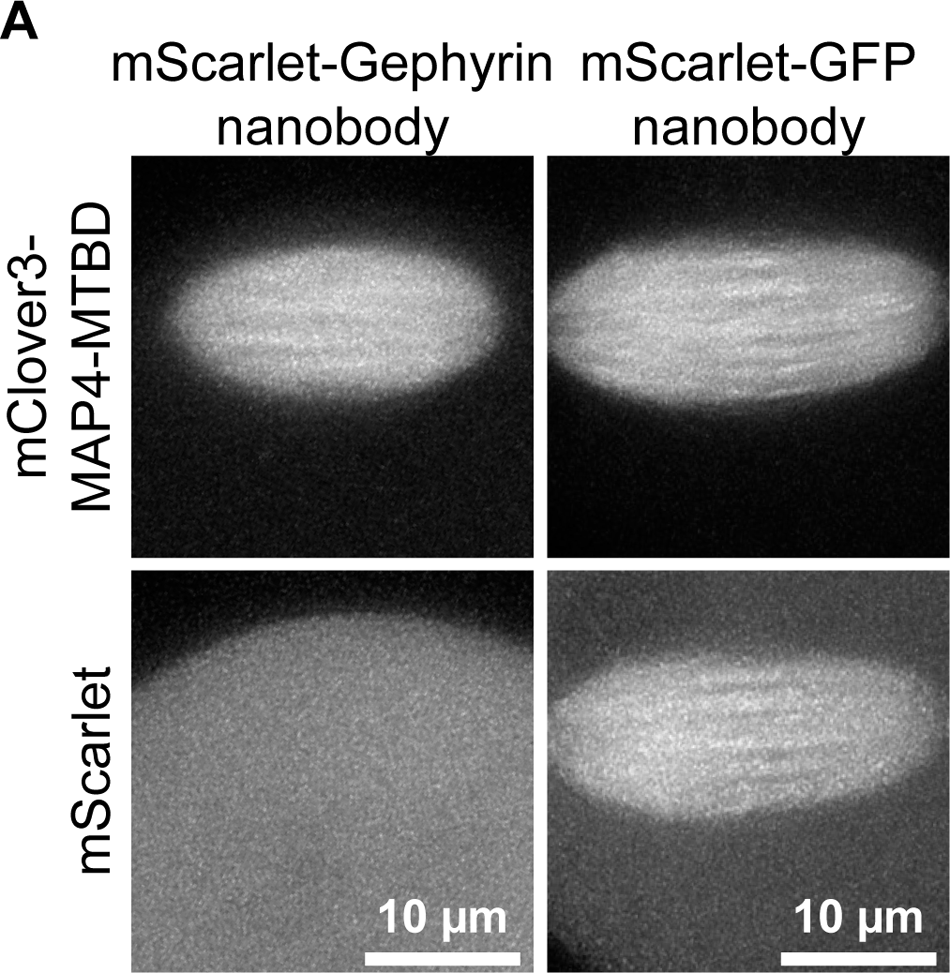
GFP-nanobodies specifically recognize mClover3 protein. A) High-resolution images of live, metaphase II-arrested wild-type mouse eggs co-expressing mClover3-MAP4-MTBD and mScarlet-tagged Gephyrin or GFP nanobodies.

**Fig. S5.**
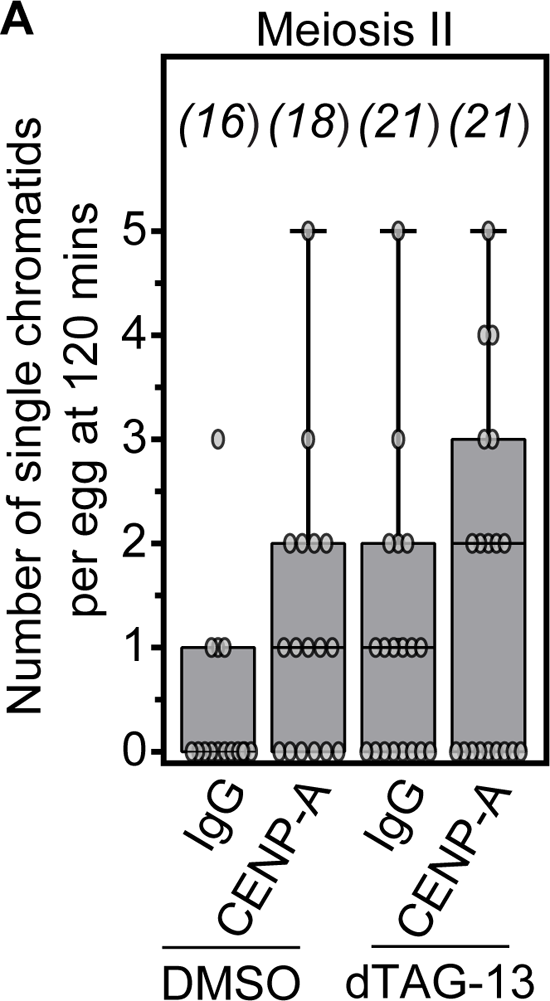
CENP-A TRIM-away increases the number of PSCS in dTAG-13-mediated cohesion degradation. (B) Quantification of the number of single chromatids per egg at 120 mins of live imaging in DMSO-treated and IgG TRIM-Away, DMSO-treated and CENP-A TRIM-Away, dTAG-13-treated and IgG TRIM-Away or dTAG-13-treated and CENP-A TRIM-Away eggs from REC8-FKBP12^F36V^-mClover3 mice.

**Table S1.**
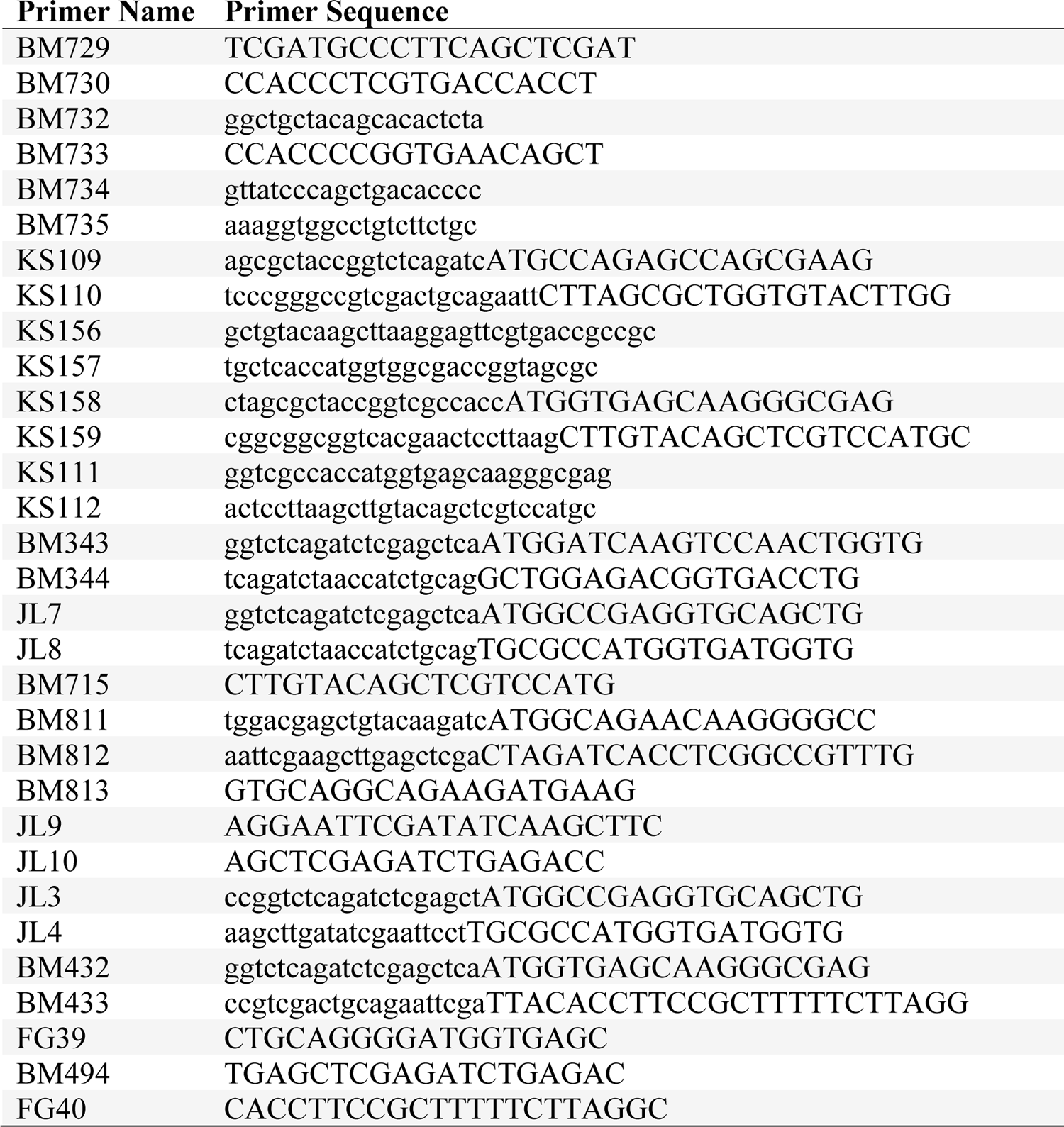
Oligonucleotide sequences used in this study.

**Movie S1.** Navigation through 1 µm apart confocal sections of REC8 (GFP antibody, grey) and homologous chromosomes (magenta) in a metaphase I-stage mouse oocyte.

**Movie S2.** High-resolution time lapse movie showing the selective removal of REC8 from chromosome arms and its retention at centromeric regions during anaphase I in a REC8-FKBP12^F36V^-mClover3 mouse oocyte.

**Movie S3.** Time lapse movie of chromosome alignment and segregation during meiosis I in a wild-type mouse oocyte. Microtubules (grey) are labeled with mNeonGreen-MAP4-MTBD and chromosomes (magenta) are labeled with H2B-mScarlet.

**Movie S4.** Time lapse movie of chromosome alignment and segregation during meiosis I in a REC8-FKBP12^F36V^-mClover3 mouse oocyte. Microtubules (grey) are labeled with mNeonGreen-MAP4-MTBD and chromosomes (magenta) are labeled with H2B-mScarlet.

**Movie S5.** Time lapse movie of chromatids (H2B-mScarlet, grey) in a DMSO-treated (dTAG-13 control), metaphase II-arrested REC8-FKBP12^F36V^-mClover3 egg.

**Movie S6.** Time lapse movie of chromatids (H2B-mScarlet, grey) in a dTAG-13-treated, metaphase II-arrested REC8-FKBP12^F36V^-mClover3 egg.

**Movie S7.** Time lapse movie of chromatids (H2B-mScarlet, grey) in a Gephyrin TRIM-Away, metaphase II-arrested REC8-FKBP12^F36V^-mClover3 egg.

**Movie S8.** Time lapse movie of chromatids (H2B-mScarlet, grey) in a GFP TRIM-Away, metaphase II-arrested REC8-FKBP12^F36V^-mClover3 egg.

**Movie S9.** Time lapse movie of chromatids (H2B-mScarlet, grey) in a DMSO-treated (CytoD control), metaphase II-arrested REC8-FKBP12^F36V^-mClover3 egg.

**Movie S10.** Time lapse movie of chromatids (H2B-mScarlet, grey) in a CytoD-treated, metaphase II-arrested REC8-FKBP12^F36V^-mClover3 egg.

**Movie S11.** Time lapse movie of chromatids (H2B-mScarlet, grey) in a dTAG-13-treated, metaphase II-arrested REC8-FKBP12^F36V^-mClover3 egg.

**Movie S12.** Time lapse movie of chromatids (H2B-mScarlet, grey) in a CytoD- and dTAG-13-treated, metaphase II-arrested REC8-FKBP12^F36V^-mClover3 egg.

**Movie S13.** Time lapse movie of chromatids (H2B-mScarlet, grey) in a DMSO-treated and IgG TRIM-Away metaphase II-arrested REC8-FKBP12^F36V^-mClover3 egg.

**Movie S14.** Time lapse movie of chromatids (H2B-mScarlet, grey) in a DMSO-treated and CENP-A TRIM-Away metaphase II-arrested REC8-FKBP12^F36V^-mClover3 egg.

**Movie S15.** Time lapse movie of chromatids (H2B-mScarlet, grey) in a dTAG-13-treated and IgG TRIM-Away metaphase II-arrested REC8-FKBP12^F36V^-mClover3 egg.

**Movie S16.** Time lapse movie of chromatids (H2B-mScarlet, grey) in a dTAG-13-treated and CENP-A TRIM-Away metaphase II-arrested REC8-FKBP12^F36V^-mClover3 egg.

